# The cellular basis of protease activated receptor type 2 (PAR2) evoked mechanical and affective pain

**DOI:** 10.1101/2020.01.31.928663

**Authors:** Shayne N Hassler, Moeno Kume, Juliet M. Mwirigi, Ayesha Ahmad, Stephanie Shiers, Andi Wangzhou, Pradipta R Ray, Serge N Belugin, Dhananjay K. Naik, Michael D. Burton, Josef Vagner, Scott Boitano, Armen N Akopian, Gregory Dussor, Theodore J Price

## Abstract

Protease-activated receptor type-2 (PAR2) has long been implicated in inflammatory and visceral pain, but the cellular basis of PAR2-evoked pain has not been delineated. While many studies have attributed PAR2-evoked pain to sensory neuron expression, RNA-sequencing experiments are ambiguous on detection of *F2rl1* mRNA. Moreover, many pharmacological tools for PAR2 have been shown to be non-specific as they also act on the Mas-related (Mrg) family of g-protein coupled receptors (GPCRs) that are highly enriched in sensory neurons. We sought to bring clarity to the cellular basis of PAR2 pain. We developed a PAR2 conditional mutant mouse by loxp targeting of exon 2 of the *F2rl1* gene and specifically deleted PAR2 in all sensory neurons using the *Pirt*^Cre^ mouse line. Our behavioral findings show that PAR2 agonist-evoked mechanical hyperalgesia and facial grimacing, but not thermal hyperalgesia, is completely dependent on PAR2 expression in sensory neurons that project to the hindpaw in male and female mice. *F2rl1* mRNA is expressed in a discrete population (~4%) of sensory neurons that also express the *Nppb* and *IL31ra* genes. This cell population has previously been implicated in itch, but our work shows that PAR2 activation in these cells causes clear pain-related behaviors from the skin. Our findings clarify the mechanism through which proteases, like tryptase and elastase, cause pain via PAR2 activation in a small subset of nociceptors.

## INTRODUCTION

Protease-activated receptors (PARs) are G-protein coupled receptors (GPCRs) that are targeted by endogenous proteases. These proteases cleave the extracellular N-terminus of the receptor to reveal a tethered peptide ligand that then induces cellular signaling (Bunnett, 2006). These receptors can signal at the cell membrane, but they also continue to generate signaling after internalization via endosomal signaling pathways (Jimenez-Vargas et al., 2018). PARs have been implicated in many different pathological states (Bao et al., 2014; Bunnett, 2006; Cocks and Moffatt, 2001; Coelho et al., 2003; Kunzelmann et al., 2002; Rothmeier and Ruf, 2012; Schaffner and Ruf, 2009). Research on PAR2 has focused on pain and inflammation due to the long-standing observation of decreased pain sensitization in *F2rl1^-/-^* mice (Vergnolle et al., 2001). Stemming from this original finding, many subsequent studies have focused on how PAR2 signaling occurs in nociceptors, but much of this body of evidence was built using a tool peptide PAR2 agonist, SLIGRL, that is now known to also be an agonist of the Mas-related GPCR, MrgprC11 (encoded by the *Mrgprx1* gene), and may also act on other Mrg receptors (Boitano et al., 2014; Liu et al., 2011). Given that the Mrg family of receptors is highly enriched in dorsal root ganglion (DRG) neurons (Meixiong and Dong, 2017; Tiwari et al., 2016), this complicates interpretation of some of the pharmacological literature on the topic. Another emerging issue in the field is that *F2rl1* gene expression in many bulk and single cell DRG sequencing datasets is either undetectable or on the threshold of detection limits (Ray et al., 2018; Usoskin et al., 2015). This includes single cell experiments from visceral afferents (Hockley et al., 2018). These findings are surprising given that PAR2 is widely considered an important therapeutic target for visceral pain with the model that PAR2 in visceral afferents is activated by endogenous proteases released during visceral inflammation (Cenac et al., 2007; Jimenez-Vargas et al., 2018; Kawabata et al., 2006; Roman et al., 2014; Zhang et al., 2011).

These disparate and somewhat controversial findings raise important questions about our understanding of PAR2 in the biology of pain. We sought to elucidate the cellular basis of PAR2-evoked pain by generating a conditional knockout mouse for the *F2rl1* gene. Importantly, another group independently generated a similar mouse and crossed it with the *Scn10a^Cre^* mouse to generate a nociceptor-specific knockout of PAR2. They found decreased mechanical hypersensitivity in response to proteases that are thought to act on PAR2, consistent with the hypothesis that PAR2 pain is mediated specifically by nociceptors (Jimenez-Vargas et al., 2018). This study also substantially advanced the field of PAR2 biology by demonstrating that PAR2 continues to signal once it is internalized via endosomal signaling. However, this study did not address different pain modalities or delineate precisely which populations of nociceptors express the *F2rl1* mRNA.

We targeted exon 2 of *F2rl1* to create a sensory neuron-specific conditional PAR2 knockout mouse using the *Pirt^Cre^* line. Our findings demonstrate that PAR2-evoked mechanical hypersensitivity and affective pain are lost in these mice while thermal hyperalgesia is lost in response to exogenous agonists but intact for endogenous protease-induced activation of PAR2. RNAscope *in situ* hybridization and cellular signaling assays on cultured mouse DRG neurons show that PAR2 is expressed by a small population of nociceptors that express several markers that identify itch nociceptors. Interestingly, we find that PAR2 activation leads only to pain, and not itch responses, demonstrating that this subpopulation of nociceptors signals pain with an appropriate stimulus.

## MATERIALS AND METHODS

### Animals

All animal protocols were approved by the University of Texas at Dallas Institutional Animal Care and Use Committee and were consistent with the NIH Guide. To generate *F2rl1^flox^* mice, lox-p sites were inserted flanking the *F2rl1* gene exon 2 on chromosome 13. The *F2rl1* gene contains 2 exons and exon 2 was targeted because it contains the majority of the coding sequence of the PAR2 protein. Mice were generated on a C57BL/6J background through a contract with Cyagen Biosciences. A neomycin selection cassette was inserted and removed through *Frt*-mediated recombination. Mice were crossed with *Pirt^Cre^* mice (Kim et al., 2016), kindly provided by Dr. Xinzhong Dong at Johns Hopkins University through a Material Transfer Agreement, at University of Texas at Dallas to generate experimental animals for behavioral experiments. Additional C57BL/6J mice were bred in our colony for cell culture and cellular anatomy studies.

The *Rosa26LSL-tDTomato*/+ mouse line on B6.129 background was obtained from the Jackson Laboratory (Bar Harbor, ME). *Trpv1^GFP^* mouse lines were purchased from the GENSAT program (MMRRC services; UNC, NC and UC Davis, CA). The *Calca^cre/+-ER^* mouse line was kindly provided by Dr. Pao-Tien Chuang (UC San Francisco, San Francisco, CA) (Song et al., 2012). Adult male mice were used in described electrophysiology experiments.

### Experimental reagents

2at-LIGRL-NH_2_ (2AT) was made as previously described (Boitano et al., 2011; Flynn et al., 2011). Neutrophil elastase (NE) was purchased from Elastin Products Company, Inc. (SE563; Owensville, Missouri). Compound 48/80 was purchased from Sigma-Aldrich (C2313).

### Behavioral methods

In behavioral experiments we used male and female mice. No sex differences were noted in any experiments so pooled data from both sexes is shown in all experiments. Behavioral observers were blinded to genotype and treatment in all experiments. Mechanical sensitivity was measured using von Frey filament testing (Chaplan et al., 1994). Animals were acclimated to suspended Plexiglas chambers (11.4 x 7.6 x 7.6 cm) with a wire mesh bottom (1 cm^2^). Withdrawal thresholds to probing the hind paws were determined prior to experimental treatment and at 1, 3, 5, 24, and 48 hours after administration. Paw withdrawal (PW) thresholds were determined by applying von Frey filaments to the non-glabrous plantar aspect of the hind paws, and a response was indicated by a withdrawal of the paw. The withdrawal thresholds were determined by the Dixon up-down method by using blinded observers. The maximum filament strength was 2 g for the experiments.

Mouse grimace scoring was performed as described by Langford et al, 2010 (Langford et al., 2010). Mice were placed individually in the same suspended Plexiglas chambers with wire mesh bottom as previously described, allowed to acclimate for 1 hour, and then scored by blinded scorers prior to experimental treatment and at 1, 3, 5, 24, and 48 hours after administration. The scores of each animal subject were averaged at each time-point by group.

Thermal sensitivity was measured using the Hargreaves method (Hargreaves et al., 1988). Mice were placed on a warmed glass floor (29°C) 20 minutes prior to each testing and using a Hargreaves apparatus (IITC Model 390), a focused beam of high-intensity light was aimed at the plantar non-glabrous surface of the hind paws. The intensity of the light was set to 30% of maximum with a cutoff value of 20 seconds. The latency to withdraw the hind paw was measured to the nearest 0.01 seconds. The hind paws were measured prior to treatment and at 1, 3, 5, 24, and 48 hours after administration.

Paw inflammation was investigated by measuring the temperature of the animal’s hind paws. All testing was performed in a climate-controlled room with an ambient temperature of 21 ± 2°C. Animals were allowed to acclimate in the testing room for 1 hour prior to testing. Colorized infrared thermograms that captured the non-glabrous surface of the animal’s hind paws were obtained using a FLIR T-Series Thermal Imaging Camera (FLIR Systems, Inc). The thermograms were captured prior to experimental treatment and at 1, 3, 5, 24, and 48 hours after administration. Thermogram analysis was performed using the FLIR Thermal Imaging Software. For each thermogram image, a straight line was drawn on the plantar surface of both hind paws and the mean temperature was recorded from the average of each pixel along the drawn line. The raw temperatures were then plotted for ipsilateral and contralateral hind paws for each individual animal.

To assess cheek itch and wipe bouts, mice were anesthetized briefly with isoflurane, and the left cheek was shaved 2-3 days prior to intradermal injection. Animals were allowed to adapt to the experimental conditions by placing them in the same suspended Plexiglas chambers with wire mesh bottom as previously described before experiments began. On the day of the experiment, mice were habituated for 1 hour in the acrylic boxes, and then their baseline behavior was recorded for 15 minutes. For each mouse, two camcorders (Samsung HMX-F90 and HMX-QF20) were placed in front of and behind the mouse, and recordings were done simultaneously. After 15 minutes of recording baseline behavior, the mice were restrained and received a 10 μl intradermal injection into the cheek of either 2AT (30 pmol, 100 pmol, or 10 nmol) or IL-31 (19 pmol). Injections were done using a Hamilton syringe (80901) with a 30G needle held parallel to the skin and inserted superficially. Once mice were injected, they were placed back into the acrylic boxes, and their behavior was video-recorded for 30 minutes. The video recordings from the two camcorders positioned in front of and behind each mouse were edited together into one video to give a simultaneous view of two different angles of the mouse. The behavior in the videos were scored by students who were blinded to the experimental groups. Wiping and scratching behaviors were scored as described previously by Shimada et al. (Shimada and LaMotte, 2008).

### DRG cultures

For primary neuronal cultures used in calcium imaging and RNAscope *in situ* hybridization, DRGs were dissected from adult ICR mice and suspended in Hanks’ Balanced Salt Solution without calcium and magnesium prior to culturing. Ganglia were incubated at 37□C for 25 minutes in 1 mg/ml papain (LS003119; Worthington), followed by 25 minutes of incubation at 37□C in 3 mg/ml collagenase type 2 (LS004176; Worthington) and 2 mg/ml Dispase II (04942078001; Sigma). Ganglia were then triturated in HBSS with a 1 ml pipette tip. The solution was passed through a 70 μm cell strainer (22363548; Fisher), and the cells were resuspended in DMEM/F12/GlutaMAX (Gibco) culture media nourished with 10% fetal bovine serum (FBS; SH30088.03; Hyclone) and 1% penicillin/streptomycin (Pen-Strep; 15070-063; Gibco). Cells were plated, allowed to adhere for 2 hours, and each well was then flooded with the same supplemented culture media described previously with additional 10 ng/ml nerve growth factor (NGF; 01-125; Millipore) and 3 ug/ml 5-fluoro-2’-deoxyuridine + 7 ug/ml uridine (FRD+U; Sigma) added. Thereafter, neurons were kept at 37□C and 5% CO_2_ in an incubator with supplemented culture media with NGF and FRD+U changed every other day until further experimentation.

DRG cultures for immunocytochemistry and RNAscope experiments were prepared as described above with the neurons plated on 8-well Chamber Slides (154534; Nunc Lab-Tek). The neurons were resuspended in culture media, plated in 100 ul in each well, and allowed to adhere for 2 hours. Then, the wells were flooded with culture media supplemented with 10% FBS, 1% Pen-Strep, 10 ng/ml NGF, and 3 ug/ml + 7 ug/ml FRD+U. Neurons were kept at 37□C and 5% CO_2_ with media changed every other day until use. Protease III (Advanced Cell Diagnostics) treatment concentration for RNAscope-ICC was optimized to 1:30.

For primary DRG neuronal cultures for electrophysiology recordings, L3-L5 DRG were quickly removed from male reporter mice *CGRP^ER-cre/+^/Rosa26^LSL-tDTomato/+^* and *CGRP^ER-cre/+^/Rosa26^LSL-tDTomato/+^/TRPV1-GFP.* DRG neurons were dissociated by treatment with a 1 mg/ml collagenase-Dispase (Roche) solution. Cells were maintained in DMEM supplemented with 2% FBS, 2 mM L-glutamine, 100 U/ml penicillin and 100 μg/ml streptomycin. The experiments were performed within 6-36 hours after DRG neuron plating.

### Calcium Imaging

DRG neurons were dissected and cultured as described before and were plated on pre-poly-D-lysine coated dishes (P35GC-1.5-10-C; MatTek) with additional laminin coating (L2020; Sigma). Neurons were used within 24 hours of plating. DRG neurons were loaded with 1 μg/μl Fura 2AM (108964-32-5; Life Technologies) for 1 hour before changing to normal bath solution (135 mM NaCl, 5 mM KCl, 10 mM HEPES, 1 M CaCl_2_, 1 M MgCl_2_ and 2 M glucose, adjusted to pH 7.4 with N-methyl-glucamine, osmolarity of 300±5 mOsm). The cells were then treated with 1 μM 2AT dissolved in normal bath solution for 120 seconds. Images were acquired on the Olympus IX73 inverted microscope at 40X magnification. For purposes of analysis, cells that responded with at least 20% ratiometric change (340 nm/380 nm) in extracellular Ca^2+^ upon treatment of KCl were classified as neurons. Out of this classification, neurons that responded with at least 40% ratiometric change upon treatment of 2AT were classified as PAR2-positive. Experiment was performed using the MetaFluor Fluorescence Ratio Imaging Software.

### Tissue preparation

Lumbar dorsal root ganglion and hind paw skin was rapidly dissected, embedded in optimal cutting temperature compound (OCT) and flash frozen immediately in dry ice. Tissues were sectioned at 20 μm onto charged slides. Sections were only briefly thawed in order to adhere to the slide but were immediately returned to the −20°C cryostat chamber until completion of sectioning.

### RNAscope in situ hybridization

RNAscope *in situ* hybridization multiplex version 1 was performed as instructed by Advanced Cell Diagnostics (ACD). Slides were transferred from the cryostat directly into cold (4°C) 10% formalin for 15 minutes and then dehydrated in 50% ethanol (5 min), 70% ethanol (5 min) and 100% ethanol (10 min) at room temperature. The slides were air dried briefly, and then boundaries were drawn around each section using a hydrophobic pen (ImmEdge PAP pen; Vector Labs). When the PAP-pen boundaries had dried, sections were incubated in protease IV reagent for 2 minutes at room temperature. Slides were washed briefly in 1X phosphate buffered saline (PBS, pH 7.4) at room temperature. Each slide was then placed in a prewarmed humidity control tray (ACD) containing dampened filter paper. For DRG experiments, *F2rl1* (PAR2; 417541; ACD), *Calca* (CGRP; 417961; ACD), and *P2rx3* (P2X3R; 521611; ACD) probes were pipetted onto each section until fully submerged and then incubated for 2 hours at 40°C. For hind paw skin, only the *F2rl1* or bacterial *dapB* (negative control) probes were used. Slides were then washed two times in 1X RNAscope wash buffer and returned to the oven for 30 minutes after submersion in AMP-1 reagent. Washes and amplification were repeated using AMP-2, AMP-3 and AMP-4 (ALT-B) reagents with a 15-min, 30-min, and 15-min incubation period, respectively. Slides were then washed two times in 0.1 M phosphate buffer (PB, pH 7.4). The DRG slides were then processed for immunohistochemistry, while the hind paw skin slides were incubated for 5 minutes in 1:5000 DAPI (ACD), washed in 0.1M PB, air dried, and cover-slipped with Prolong Gold.

### Immunohistochemistry

After completion of RNAscope *in situ* hybridization, DRG slides were incubated in blocking buffer (10% Normal Goat Serum, 0.3% Triton-X 100 in 0.1 M PB) for 1 hour at room temperature while being shielded from light. Slides were placed in a light-protected humidity-controlled tray and incubated in primary antibody (mouse-anti-Neurofilament 200; clone N52; MAB5266; Sigma) at 1:500 in blocking buffer overnight at 4°C. The next day, slides were washed two times in 0.1 M PB and then incubated in secondary antibody (goat-anti-mouse IgG (H+L) Alexa Fluor 405; 1:2000; A-31553; Invitrogen) for 1 hour at room temperature. Sections were washed two times in 0.1 M PB, air-dried, and cover-slipped with Prolong Gold Antifade mounting medium.

### Image Analysis for DRG sections

Three mice per genotype were imaged on an Olympus FV3000 confocal microscope at 20X magnification. One image was acquired of each mouse DRG section, and 3 sections were imaged per mouse (total: 9 images). The raw image files were brightened and contrasted equally in Olympus CellSens software and then analyzed manually one cell at a time for expression of *Calca, P2rx3*, and *F2rl1.* Cell diameters were measured using the polyline tool. The combination of NF200 signal (not shown), *Calca, P2rx3*, and *F2rl1* were used to quantify the total neuronal population. Representative images of hind paw skin are shown from *F2rl1^flox^Pirt*^+/+^ and *F2rl1^flox^Pirt^Cre^* mice with a negative control from a *F2rl1^flox^Pirt*^+/+^ animal imaged at the same settings.

### RNAscope in situ hybridization on DRG cultures

DRG cultures were prepared as described with the neurons plated on 8-well Chamber Slides (154534; Nunc Lab-Tek) coated with poly-D-lysine (P0899; Sigma-Aldrich). On day 5, the cultures were treated with 1 μM 2AT or vehicle (culture media) for 10 minutes in the incubator. The samples were then prepared as instructed by ACD. The chambers were disassembled, and the slides submerged in 1X PBS. They were transferred to 10% formalin for 30 minutes at room temperature, followed by three washes in 1X PBS. Hydrophobic boundaries were drawn around each well as previously described. Each well was incubated with protease III reagent (1:30 in 1X PBS) for 10 minutes at room temperature. Slides were washed in 1X PBS and then placed in a prewarmed humidity control tray containing dampened filter paper. *F2rl1* and *P2rx3* probes were pipetted onto each well. Two wells received only control probes, negative (bacterial *dapB*) or positive (320881; ACD). Slides were incubated in the probes, followed by washes and amplification as previously described. After completion of RNAscope *in situ* hybridization, immunocytochemistry was performed.

### Immunocytochemistry

The following steps were performed in a light-protected humidity control tray. Slides were incubated in blocking buffer (10% Normal Goat Serum in 0.1 M PB) with 0.02% Triton-X 100 for 1 hour at room temperature. They were then incubated overnight at 4°C with primary antibody, mouse-anti-Neurofilament 200 (NF200; clone N52; MAB5266; Sigma) at 1:500 and rabbit-anti-P-p44/42 MAPK T202/Y204 (p-ERK; 9101; Cell Signaling Technology) at 1:250 in blocking buffer. The next day, slides were washed twice in 0.1 M PB and incubated for 1 hour at room temperature with secondary antibody, goat-anti-mouse IgG (H+L) Alexa Fluor 405 (A-31553; Invitrogen) and goat-anti-rabbit IgG (H+L) Alexa Fluor 647 (A-21245; Invitrogen), both at 1:2000 in blocking buffer. Slides were washed twice in 0.1 M PB, air-dried, and cover-slipped with Prolong Gold Antifade mounting medium.

### Image Analysis for DRG cultures

5 wells of each treatment were imaged using an Olympus confocal microscope (FV1200). 3-6 images were taken per well at 40X magnification with 9 z-slices for a total of 22 images per treatment. The raw image files were projected to their maximum z, brightened and contrasted equally in Olympus CellSens software, and analyzed manually for expression of *P2rx3* and *F2rl1.* NF200 signal was used to verify the neuronal population. Z slices that did not contain the neuron in focus were excluded from analysis. ROIs of each neuron were drawn using the ellipse tool, and p-ERK signal was quantified using mean gray intensity value. Background values taken from the negative control were subtracted prior to analysis. Percentage of neurons expressing *P2rx3* and *F2rl1* were summed from both treatment groups, while p-ERK intensity was compared among 2AT and vehicle treated *F2rl1* positive and negative neurons. Representative images at 40X are shown for *P2rx3, F2rl1* and NF200 expression, along with a zoom-in of a single neuron. Zoom-in of a single neuron is also shown for p-ERK expression in a representative *F2rl1* positive neuron.

### Electrophysiology

Recordings were made in whole-cell current clamp configurations at 22-24°C. Data were acquired and analyzed using an Axopatch 200B amplifier and pCLAMP10.2 software (Molecular Devices, Sunnyvale, CA). Recorded data were filtered at 5 kHz and sampled at 30 kHz. Borosilicate pipettes (Sutter, Novato, CA) were polished to resistances of 2-3 MΩ. Access resistance (R_s_) was compensated (40-80%) when appropriate up to the value of 6-8 MΩ. Data were rejected when R_s_ changed >20% during recording, leak currents were >50 pA, or input resistance was <300 MΩ. Standard external solution (SES) contained (in mM): 140 NaCl, 5 KCl, 2 CaCl_2_, 1 MgCl_2_, 10 D-glucose, and 10 HEPES, pH 7.4. The standard pipette (internal) solution (SIS) contained (in mM): 140 KCl, 1 MgCl_2_, 1 CaCl_2_, 10 EGTA, 10 D-glucose, 10 HEPES, pH 7.3, 2.5 ATP, and 0.2 GTP. Drugs were applied by a fast, pressure-driven and computer-controlled 4-channel system (ValveLink8; AutoMate Scientific, San Francisco, CA) with quartz application pipettes.

CGRP^+^ DRG small (<30 pF) neurons from *CGRP^cre/+-ER^/Rosa26^LSL-tDTomato/+^* mice were randomly selected for recording. TRPV1^+^/CGRP^-^ DRG neurons from *CGRP^cre/+-ER^/Rosa26^LSL-tDTomato/+^/TRPV1-GFP* reporter mice were selected for recording as they have PAR2 (Usoskin et al., 2015). To characterize modulation of TRPV1^+^/CGRP^-^ or CGRP^+^ DRG neuron excitation by vehicle (control) or PAR2 activating peptide (2AT), the following sequence of recording protocols were applied: (1) a single AP in current clamp configuration was generated with a 0.5 ms and 1 nA current step to define the type of sensory neurons (Patil et al., 2018); (2) a linear ramp from 0 to 0.1 nA for 1 sec was applied to generate a control AP train; (3) the patched neuron was treated for 2-5 min with vehicle or PAR2 activator; and then (4) the ramp as in the step 2 was re-applied. Data was accumulated from 3-5 independent mouse DRG neuronal cultures. Each culture was generated from one male mouse. Changes in neuronal excitability were calculated by dividing AP frequency generated by a current ramp after vehicle or drug-treatment to AP frequency produced by the ramp before treatment. Excitability was determined to be regulated by 2AT when the drug treatment produced statistically significant increase in AP frequency than vehicle-treatment (i.e. control).

### Bioinformatics

Read counts for each coding gene for 204 single cell RNA-sequencing profiles of mouse DRG sensory neurons were obtained from Gene Express Omnibus deposit (accession number GSE63576) (Li et al., 2016). t-SNE based clustering and visualization of the single cell data sets was performed using Seurat package 2.2.1 (Butler et al., 2018; Gribov et al., 2010).

### Data Analyses and Statistics

All statistical tests done using GraphPad Prism version 8.3.0 (GraphPad Software, Inc. San Diego, CA USA). Differences between groups were evaluated using one- and two-way ANOVAs followed by either Bonferroni’s, Tukey’s, Dunnett’s, or Sidak’s multiple comparisons for data sets with three or more groups. Unpaired t-tests were done for data sets with only two groups as indicated in the text and figure captions. Outliers were assessed using a Grubb’s test and excluded. Only 1 outlier datapoint was identified in this study and is noted in the figure caption for that dataset. All statistics including t, q, DF and exact p-values are shown in supplementary tables. All data are represented as mean ± SEM with p < 0.05 considered significant.

## RESULTS

### Evaluation of PAR2 expression in DRG sensory neurons

Recent RNA-sequencing studies find very low expression levels for *F2rl1* mRNA in DRG (Hockley et al., 2018; Ray et al., 2018; Usoskin et al., 2015), a surprising finding given the large literature on PAR2 signaling in DRG neurons (Bunnett, 2006). We re-evaluated *F2rl1* mRNA expression in a deeply sequenced single-cell RNA-sequencing dataset (Li et al., 2016). We found that *F2rl1* mRNA was detected, but only in a small subset of cells (Fig 1). Single-cell expression for *F2rl1* was identified in a subpopulation of neurons with gene markers *Il31ra*, and *Nppb*, that co-express *Hrh1*, and *Mrgprx1*, all genes thought to mark a set of sensory neurons that are important for itch sensation (Lamotte et al., 2013; Meixiong and Dong, 2017; Mishra and Hoon, 2013). These neurons also co-expressed *Trpv1.*

**Figure 1.**
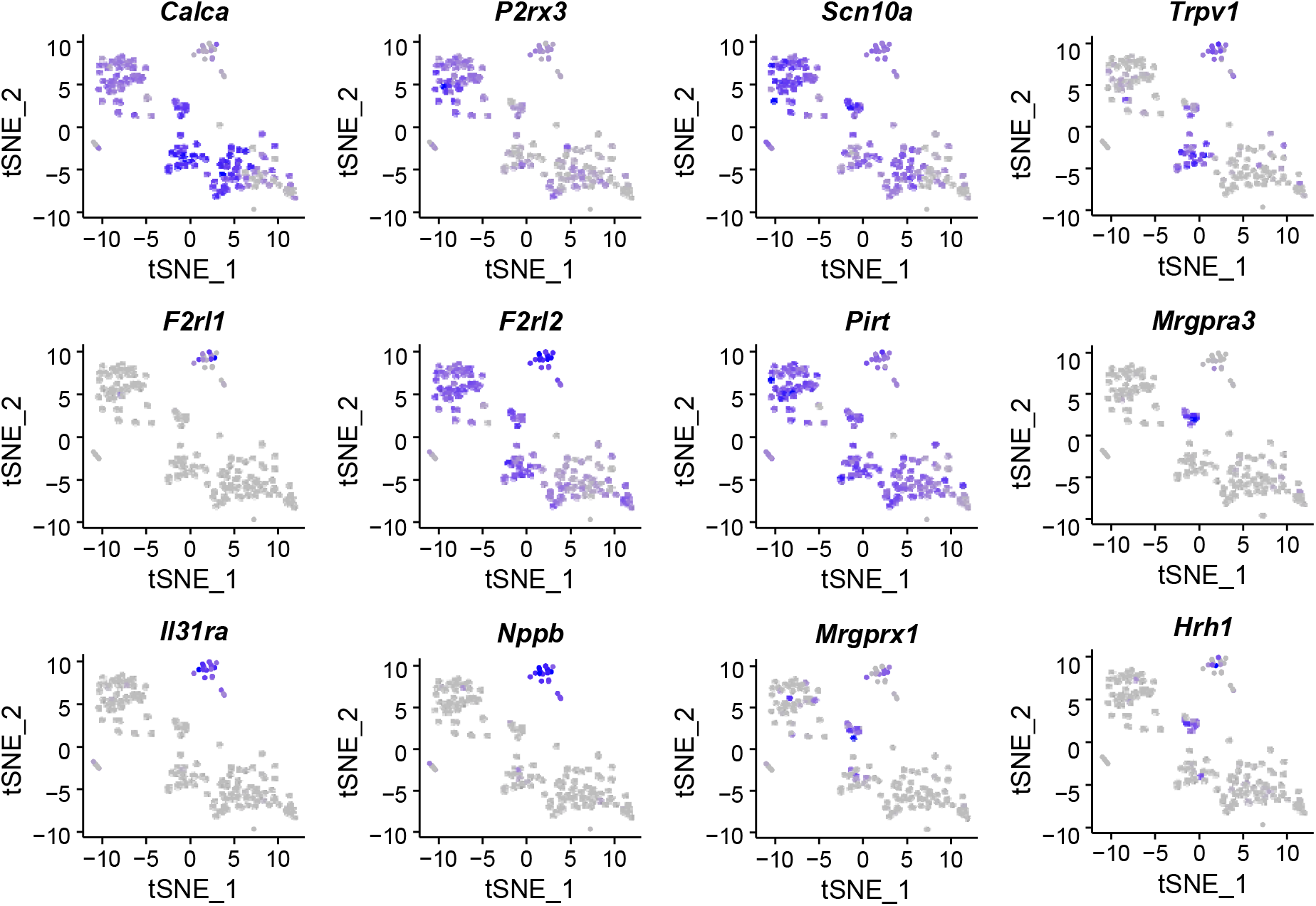
Delineation of *F2rl1* expressing DRG neurons from single cell RNA sequencing experiments. Single-cell RNA sequencing of mouse DRG neurons demonstrates that *F2rl1* mRNA is expressed in a discrete population of sensory neurons that also express the gene markers *Nppb* and *Il31ra.*

We next evaluated *F2rl1* mRNA expression in mouse DRG using RNAscope. We used probes for mRNAs encoding PAR2, CGRP, and P2X3R in triple-labeling experiments (Fig 2A). PAR2 expression was found in a small subset (3.4%) of neurons in WT mice. We generated a conditional knockout of PAR2, *F2rl1^flox^Pirt^Cre^*, and tested for knockout of PAR2 mRNA expression in DRG of WT, *F2rl1^flox^Pirt^+/+^*, and *F2rl1^flox^Pirt^Cre^* mice. We detected PAR2 mRNA expression in WT mice but not conditional knockout mice (Fig 2B). In WT mice, PAR2 expression was rare in cells expressing mRNA encoding CGRP, but most PAR2 mRNA-expressing cells also co-expressed P2X3R (Fig 2C-E). PAR2 mRNA-expressing cells were almost entirely small-diameter (Fig 2D). As an additional control for specificity of our conditional knockout approach, we did RNAscope experiments on skin sections from *F2rl1^flox^Pirt^+/+^* and *F2rl1^flox^Pirt^Cre^* mice. We observed PAR2 mRNA expression in populations of skin cells in both genotypes, demonstrating that PAR2 is not knocked out in skin cells using the *Pirt^Cre^* approach (Fig 2F).

**Figure 2.**
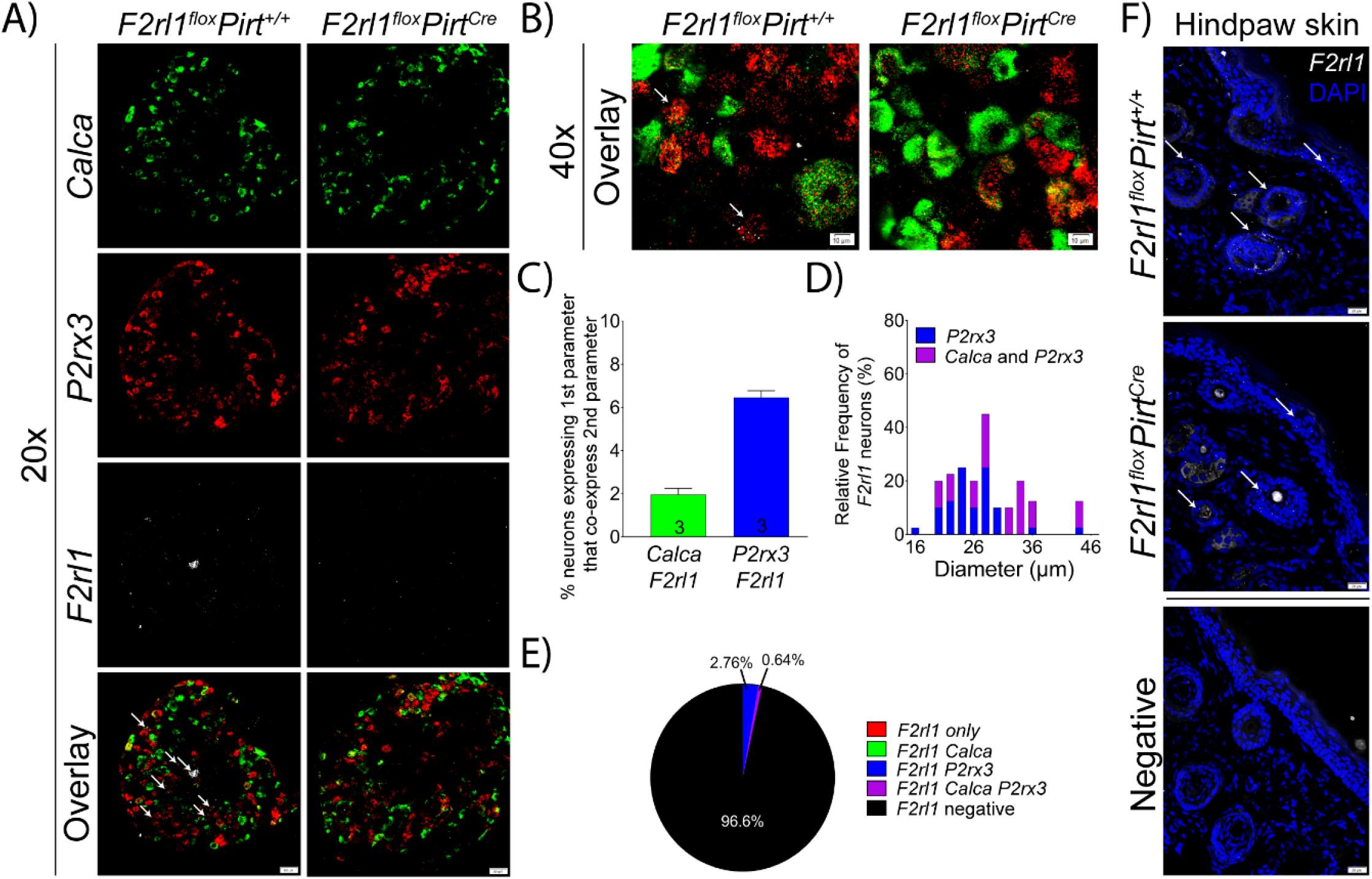
*F2rl1* is expressed by a small subset of sensory neurons. A) Representative 20X images of *Calca*, *P2rx3*, and *F2rl1* mRNA signal in the DRG of *F2rl1^flox^Pirt^+/+^* and *F2rl1^flox^Pirt^cre^* mice. B) 40X overlay image showing RNAscope signal at the cellular level. C) Percentage of *Calca-* and *P2rx3*-positive neurons that co-express *F2rl1* D) Histogram illustrating the diameter of neurons expressing *F2rl1.* E) Pie chart illustration of the percentage of *F2rl1*-positive cells that colocalize with *Calca- and P2rx3*-positive neurons. F) Representative hind paw skin images from *F2rl1^flox^Pirt^+/+^* and *F2rl1^flox^Pirt^cre^* mice with a negative probe control show the specificity of conditional knockout of *Frl1* expression in sensory neurons and not skin cells.

### PAR2 agonist-induced signaling occurs exclusively in PAR2-expressing DRG neurons

Having established that PAR2 mRNA expression is restricted to a small proportion of DRG neurons, we tested whether this restricted expression pattern would be found in functional assays. We began with Ca^2+^ imaging of DRG neurons in culture. We found that the specific PAR2 agonist, 2AT (1 μM (Boitano et al., 2011; Flynn et al., 2011)), induced Ca^2+^ signaling in ~4% of DRG neurons (Fig 3A-C). We also assessed whether PAR2-mediated plasticity could be observed in DRG neurons using patch clamp electrophysiology. To do this, we focused on *Trpv1*-expressing neurons using a genetically-tagged line because our analysis of RNA-sequencing data revealed that PAR2 overlaps with a subset of *Trpv1*-expressing cells. As predicted, we found that ramp-evoked spiking was augmented by 2AT (1 μM, 3 min) treatment but only in CGRP-negative/TRPV1-positive cells (Fig 3D-E).

**Figure 3.**
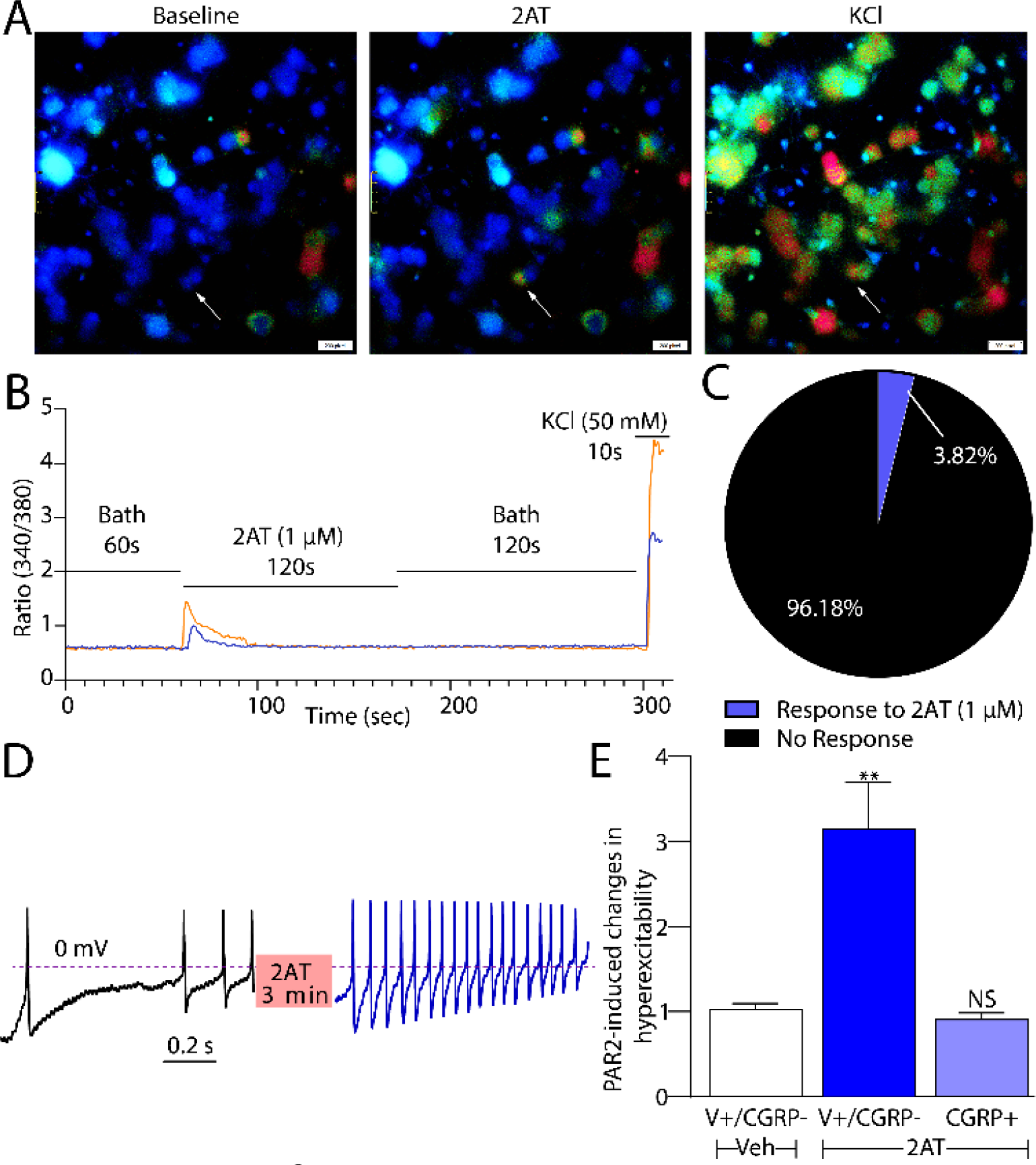
2AT-evoked Ca^2+^ signaling is specific for PAR2-expressing neurons. A) Representative images of cultured DRG neurons at baseline and upon treatment with 2AT (1 μM) and KCl (50 mM), a positive control for neuronal Ca^2+^ signaling. B) Representative traces of cultured DRG neurons with a 340/380 ratiometric change during Ca^2+^ imaging. C) Pie chart illustrating the percentage of PAR2-positive neurons in culture as characterized by response to 2AT (1 μM). D) Whole-cell current clamp recordings reveal increased firing of TRPV1^+^/CGRP^-^ cultured DRG neurons after activation with 2AT (1 μM). E) Electrophysiological experiments demonstrate that 2AT (1 μM) induced hyperexcitability exclusively in TRPV1^+^/CGRP^-^ neurons. n = 7 for TRPV1^+^/CGRP^-^ neurons treated with vehicle, n = 13 for TRPV1^+^/CGRP^-^ neurons treated with 2AT (1 μM), and n = 8 for CGRP^+^ neurons treated with 2AT (1 μM). One-way ANOVA with Tukey’s multiple comparisons (Panel E) **p<0.01.

PAR2 agonists can cause development of chronic pain via activation of extracellular signal-regulated protein kinase (ERK1/2) signaling (Tillu et al., 2015). We also tested whether 2AT evokes activation of ERK signaling specifically in PAR2 mRNA expressing cells. First, we evaluated PAR2 mRNA expression in DRG cultures from mice using RNAscope. We observed PAR2 expression in ~3% of cells, almost all of which also expressed P2X3R mRNA (Fig 4 A-B). We then exposed cultured mouse DRG neurons to 2AT (1 μM, 10 min) and then did RNAscope for PAR2 mRNA and ICC for p-ERK. Strikingly, we observed increased ERK phosphorylation but only in cells that also expressed PAR2 mRNA (Fig 4C-D). These experiments demonstrate that 2AT acts specifically on PAR2-expressing cells to induce increased intracellular Ca^2+^, augmented cellular excitability, and enhanced ERK activity.

**Figure 4.**
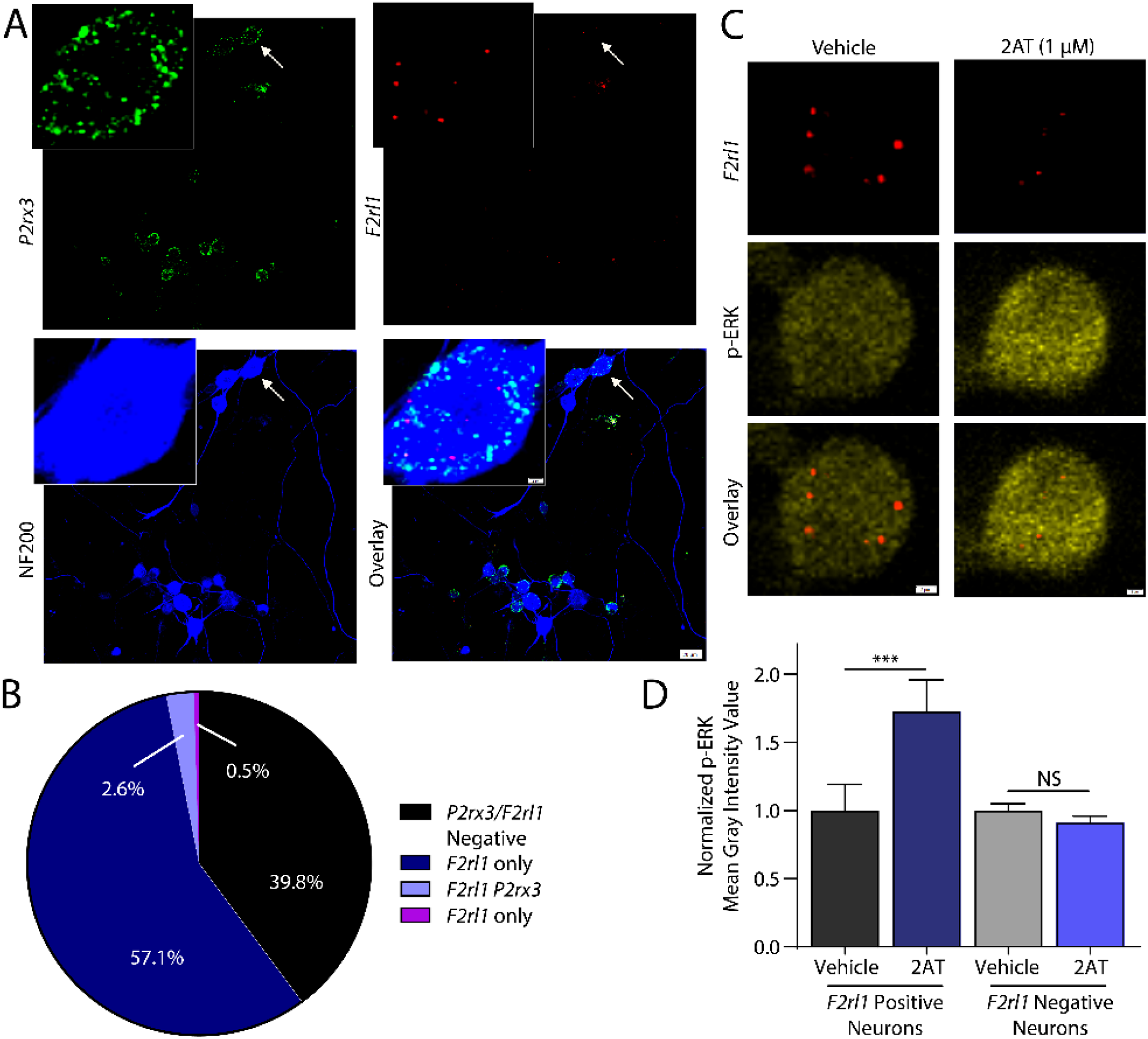
2AT-evoked p-ERK signaling is specific for *F2rl1*-expressing neurons. A) Representative images of *P2rx3, F2rl1* mRNA, and NF200 protein signal of cultured DRG neurons from WT mice. Smaller image panels display zoom-in of a single neuron. B) Pie chart illustrating the percent distribution of neuronal *F2rl1* and *P2rx3* expression *in vitro.* C) Representative images of *F2rl1* mRNA signal and p-ERK immunolabeling in cultured DRG from WT mice after treatment with vehicle or 2AT (1 μM). D) Signal intensity of p-ERK increases markedly in *F2rl1*-positive neurons after treatment with 2AT (1 μM). n = 15 and n = 16 for *F2rl1*-positive neurons treated with vehicle or 2AT, respectively. n = 88 for both vehicle and 2AT treatment groups in *F2rl1*-negative neurons. One-way ANOVA with Bonferroni’s multiple comparisons (Panel D) ***p<0.001.

### PAR2-driven mechanical hyperalgesia and grimace are mediated by sensory neurons

In order to determine if PAR2 receptor activity in DRG sensory neurons is responsible for specific types of nociceptive behaviors, we injected either the PAR2 agonist, 2AT, the mast cell degranulator 48/80, or neutrophil elastase (NE) into the hind paws of either control *F2rl1^flox^Pirt^+/+^* or PAR2 conditional knockout *F2rl1^flox^Pirt^Cre^* mice. Using von Frey testing, we found that when 2AT was injected into the hind paws of mice lacking PAR2 in sensory neurons (*F2rl1^flox^Pirt^Cre^)*, the mice showed only a very transient mechanical hypersensitivity (Fig 5A). In contrast, *F2rl1^flox^Pirt^+/+^* mice displayed mechanical hypersensitivity that lasted for at least 24 hours and that was significantly greater than PAR2 conditional knockout mice (Fig 5A). Strikingly similar results were obtained for both 48/80 (Fig 5B) and NE (Fig 5C), demonstrating that PAR2-mediated mechanical hypersensitivity requires sensory neuron PAR2 expression in mice.

**Figure 5.**
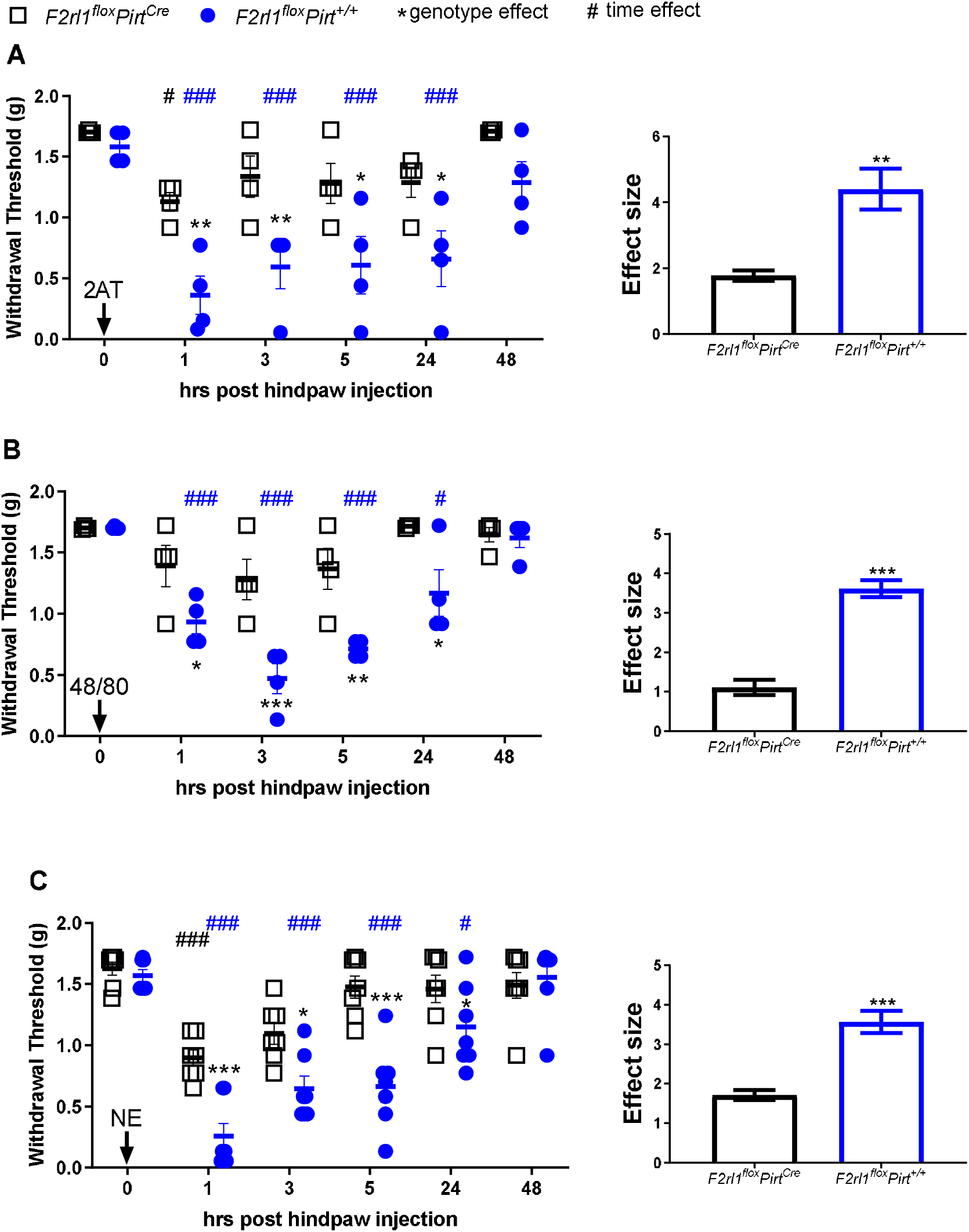
PAR2-induced mechanical hypersensitivity is sensory neuron mediated. Mice were injected with PAR2 agonists before assessing mechanical sensitivity. 2AT (30 pmol) panel A, 48/80 (6.5 nmol) panel B, and NE (10 units) panel C. *p < 0.05 compared with *F2rl1^flox^Pirt^+/+^* or *F2rl1^flox^Pirt^Cre^ groups.* #p < 0.05 compared with baseline measures. n = 4 for *F2rl1^flox^Pirt^+/+^* and *F2rl1^flox^Pirt^Cre^* groups treated with 2AT and 48/80, n = 7 for *F2rl1^flox^Pirt^+/+^* and *F2rl1^flox^Pirt^Cre^* groups treated with NE. Two-way ANOVA with Sidak’s and Dunnett’s multiple comparisons (Panel A-C) *p<0.05, **p<0.01, ***p<0.001. Unpaired t-test **p<0.01, ***p<0.001.

To determine if PAR2 activation results in changes in affective measures of pain, we injected 2AT, 48/80, and NE into the hind paws of either *F2rl1^flox^Pirt^/+^* or *F2rl1^flox^Pirt^Cre^* mice and recorded grimacing behaviors. *F2rl1^flox^Pirt^Cre^* did not show any signs of grimacing in response to 2AT injection while *F2rl1^flox^Pirt^+/+^* showed a significant increase in mouse grimace scores for up to 5 hours after injection (Fig 6A). 48/80 (Fig 6B) and NE (Fig 6C) likewise caused grimacing for about 5 hours in *F2rl1^flox^Pirt^+/+^*, but no grimacing responses were noted in *F2rl1^flox^Pirt^Cre^* mice. Therefore, grimacing in response to PAR2 activation also depends on PAR2 expression in DRG neurons.

**Figure 6.**
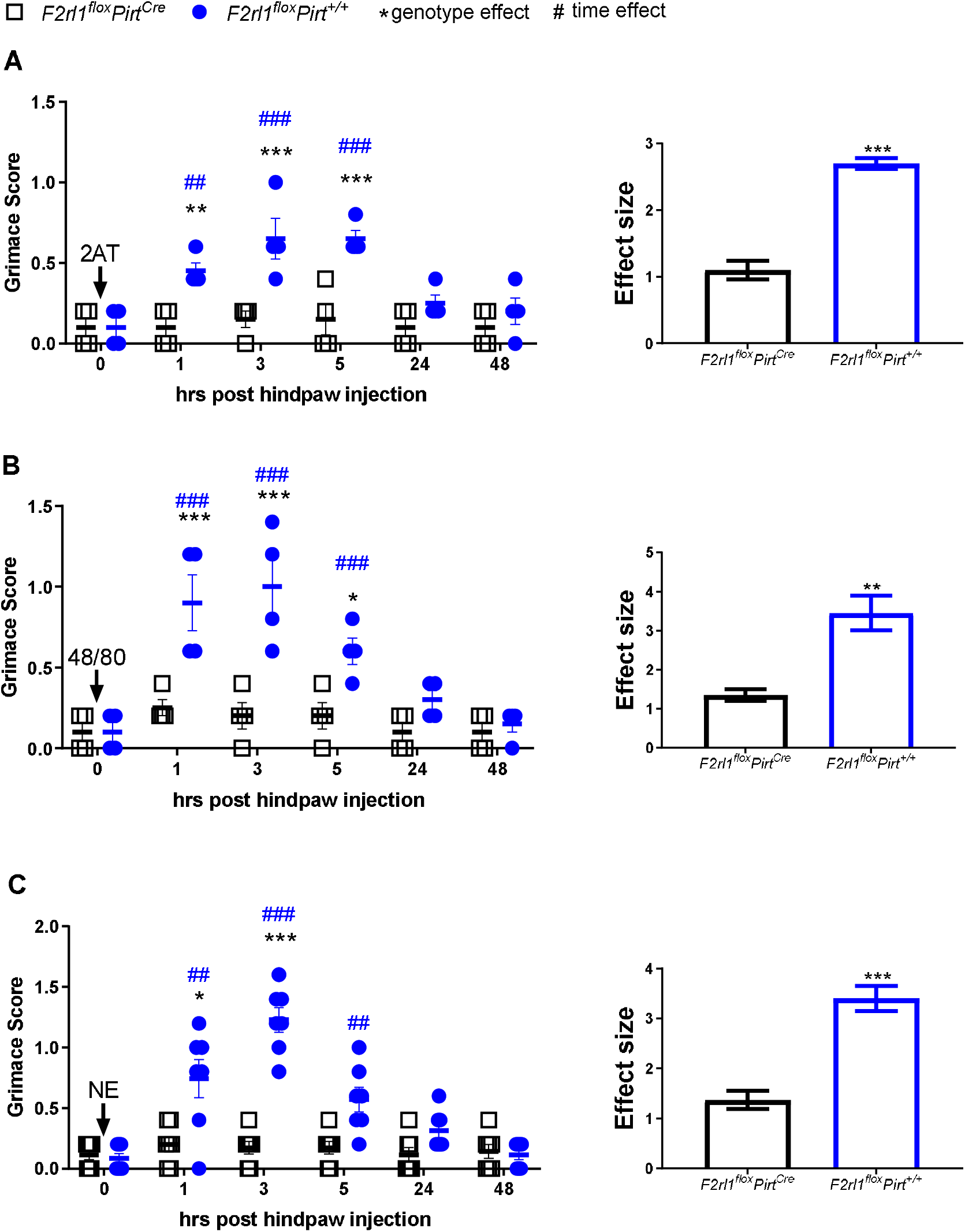
PAR2 agonists effectuate facial grimacing via sensory neurons. Mice were injected with PAR2 agonists and grimacing was subsequently scored. 2AT (30 pmols) panel A, 48/80 (6.5 nmols) panel B, and NE (10 units) panel C. *p < 0.05 compared with *F2rl1^flox^Pirt^+/+^* or *F2rl1^flox^Pirt^Cre^* groups. ^#^p < 0.05 compared with baseline measures. n = 4 for *F2rl1^flox^Pirt^+/+^* and *F2rl1^flox^Pirt^Cre^* groups treated with 2AT and 48/80, n = 7 for *F2rl1 ^flox^Pirt^+/+^* and *F2rl1^flox^Pirt^Cre^* groups treated with NE. Two-way ANOVA with Sidak’s and Dunnett’s multiple comparisons (Panel A-C) *p<0.05, **p<0.001, ***p<0.01. Unpaired t-test **p<0.01, ***p<0.001.

Thermal hyperalgesia findings were less clear-cut than mechanical sensitivity and grimace. 2AT injected into the hind paws of *F2rl1^flox^Pirt^Cre^* did not cause thermal hyperalgesia when compared to baseline (Fig 7A). *F2rl1^flox^Pirt^+/+^* mice showed thermal hyperalgesia only at the 24-hour time point, and the effect size of thermal hyperalgesia was greater in mice with intact PAR2 expression in DRG neurons (Fig 7A). 48/80 caused significant thermal hyperalgesia at the 5- and 48-hour time points only in *F2rl1^flox^Pirt^Cre^* mice, but the effect size did not differ between genotypes (Fig 7B). NE caused robust thermal hyperalgesia in both genotypes that lasted for 48 hours (Fig 7C).

**Figure 7.**
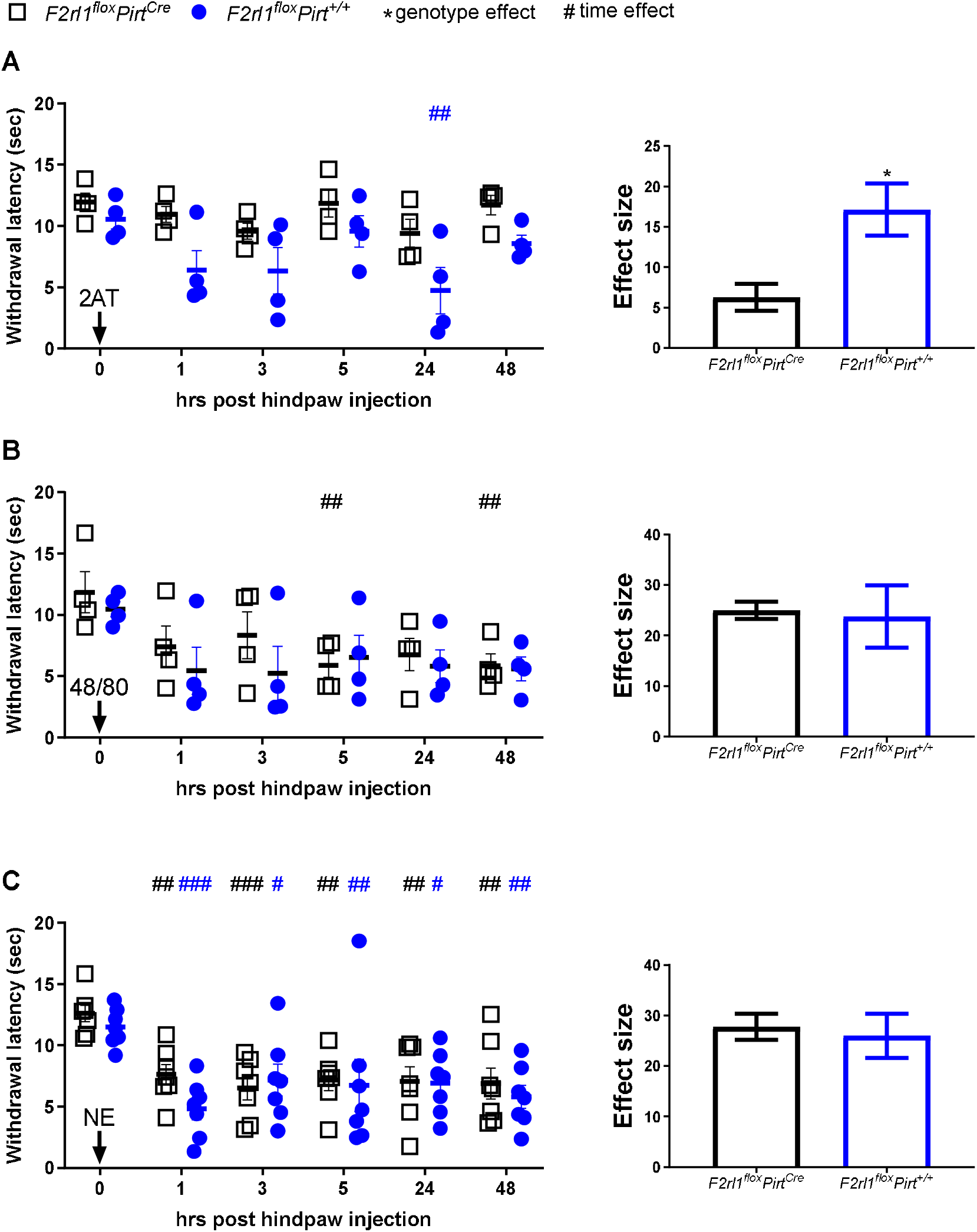
Effects of 2AT, 48/80, and NE on thermal hyperalgesia. Mice were injected with PAR2 agonists and then latency to paw withdrawal was measured. 2AT (30 pmols) panel A, 48/80 (6.5 nmols) panel B, and NE (10 units) panel C. *p < 0.05 compared with *F2rl1^flox^Pirt^+/+^* or *F2rl1^flox^Pirt^Cre^* groups. ^#^p < 0.05 compared with baseline measures. n = 4 for *F2rl1^flox^Pirt^+/+^* and *F2rl1^flox^Pirt^Cre^* groups treated with 2AT and 48/80, n = 7 for *F2rl1^flox^Pirt^+/+^* and *F2rl1^flox^Pirt^Cre^* groups treated with NE. Two-way ANOVA with Sidak’s and Dunnett’s multiple comparisons (Panel A-C) *p<0.05, **p<0.01, ***p<0.001. Unpaired t-test *p<0.05.

To determine if PAR2 receptor activation results in changes in temperature indicative of inflammation of the paw, we used infrared FLIR imaging. With 2AT injection, we did not note any change in hind paw temperature in either genotype at any time point (Fig 8A). On the other hand, 48/80 caused a significant increase in paw temperature but only in *F2rl1^flox^Pirt^+/+^* mice (Fig 8B). Upon injection of NE into the hind paw, we again observed that only the *F2rl1^flox^Pirt^+/+^* showed a significant increase in paw temperature (Fig 8C).

**Figure 8.**
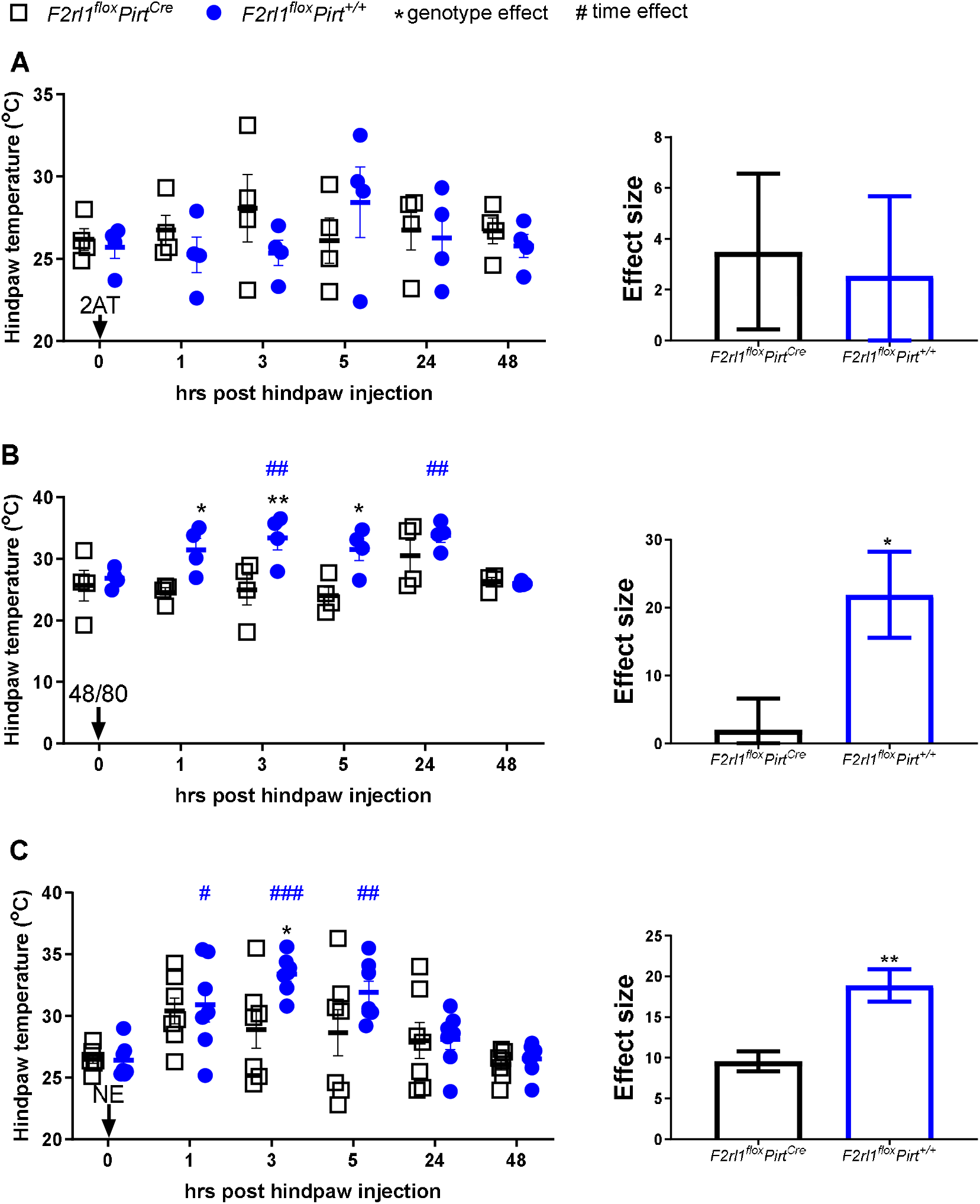
Effects of 2AT, 48/80, and NE on paw temperature are sensory neuron PAR2-mediated. Mice were injected with PAR2 agonists and then hind paw temperatures were measured. 2AT (30 pmol) panel A, 48/80 (6.5 nmol) panel B, and NE (10 units) panel C. *p < 0.05 compared with *F2rl1^flox^Pirt^+/+^* or *F2rl1^flox^Pirt^Cre^* groups. ^#^p < 0.05 compared with baseline measures. n = 4 for *F2rl1^flox^Pirt^+/+^* and *F2rl1^flox^Pirt^Cre^* groups treated with 2AT and 48/80, n = 7 for *F2rl1^flox^Pirt^+/+^* and *F2rl1^flox^Pirt^Cre^* groups treated with NE. Two-way ANOVA with Sidak’s and Dunnett’s multiple comparisons (Panel A-C) *p<0.05, **p<0.01, ***p<0.001. Unpaired t-test *p<0.05, **p<0.01.

Our data shows that *F2rl1* mRNA is very specifically expressed in a subset of sensory neurons that are associated with itch behavior. Therefore, we assessed acute itch and pain behaviors using the cheek scratch vs. wipe assay (Shimada and LaMotte, 2008). We used 3 concentrations of 2AT, 2 of which should be specific for PAR2 and a higher concentration that could potentially activate Mrg receptors (Boitano et al., 2014; Liu et al., 2011). We also used interleukin-31 (IL-31, 19 pmol) since the PAR2-expressing population of cells expresses the IL-31 receptor (IL-31R), and IL-31R signaling is linked to itch behaviors (Cevikbas et al., 2014). We observed that IL-31 caused an increase in itch bouts, but low doses of 2AT (30 and 100 pmol) did not (Fig 9A). The higher dose of 2AT (10 nmol) did cause itch bouts in WT mice, but this effect was not seen in global *F2rl1^-/-^* mice. IL-31 and both low doses of 2AT caused wipes, indicative of pain behaviors, in WT mice (Fig 9B). The higher dose of 2AT did not cause significant wiping behavior in WT mice but did cause wiping in *F2rl1^-/-^* mice. These findings show that 2AT, at concentrations that are specific for PAR2 activation (Boitano et al., 2014), only causes acute pain behaviors and not itching.

**Figure 9.**
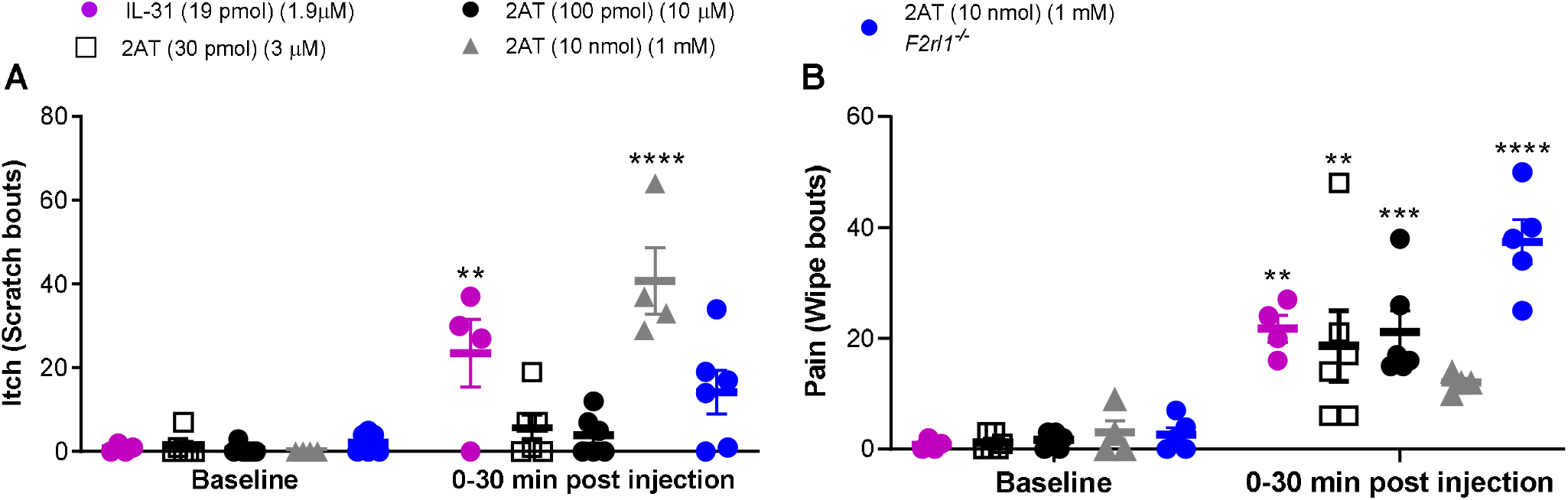
Low-dose administration of 2AT induces pain, but not itch bouts. Intradermal injections of IL-31 (19 pmol) and 2AT (30 pmol, 100 pmol and 10 nmol) were administered on the left cheek. Itch (scratch bouts) and pain (wipe bouts) were scored up to 30 minutes after injections (Panel A and B). n = 4, n = 6, n = 6, and n = 4 for WT mice treated with IL-31 (30 pmol) and 2AT (30 pmol, 100 pmol and 10 nmol), respectively, and n = 6 for *F2rl1^-/-^* mice treated with 2AT (10 nmol). Two-way ANOVA with Bonferroni’s multiple comparisons (baseline versus treatment) **p<0.01, ***p<0.001, ****p<0.0001

## DISCUSSION

PAR2 was one of the first pain targets identified using knockout mouse technology (Vergnolle et al., 2001) and has remained a prominent target in the pain field for 2 decades (Bao et al., 2014; Bunnett, 2006). While many aspects of PAR2 physiology and pharmacology have been revealed, the elucidation of other targets, such as the Mrg family of GPCRs for some broadly used PAR2 tools has complicated interpretation of much of the existing literature (Liu et al., 2011). Moreover, several recent RNA sequencing papers have reported surprisingly low levels of *F2rl1* gene expression in DRG and/or in single DRG neurons (Hockley et al., 2018; Li et al., 2016; Ray et al., 2018). Our experiments were aimed at gaining better clarity on which neurons in the DRG express PAR2 and what aspects of pain behavior are driven by these neurons. Combined with a recent publication that independently generated a nociceptor-specific *F2rl1* knockout mouse with nearly identical findings (Jimenez-Vargas et al., 2018), our work makes it clear that sensory neuron-expressed PAR2 is required for mechanical hypersensitivity and spontaneous pain behaviors caused by PAR2 activation in the paw. A unique aspect of our work is that we demonstrate that this effect is driven by a very small population of DRG neurons. We conclude that sensory neuron-expressed PAR2 is a key target for mechanical and spontaneous pain driven by the release of endogenous proteases from many types of immune cells.

Thermal hyperalgesia caused by inflammation is at least partially mediated by PAR2 (Vergnolle et al., 2001). In our experiments we did not observe any genotype differences in 48/80- or NE-evoked thermal hyperalgesia, indicating that this effect is likely driven by PAR2 expression in non-neuronal cells. Interestingly, PAR2 is expressed by a small subset of DRG nociceptors that also express TRPV1, but not CGRP, a receptor that is required for the generation of thermal hyperalgesia in inflammatory conditions (Caterina et al., 2000; Caterina et al., 1997; Tominaga et al., 1998). It is possible that PAR2 is not expressed by a sufficient proportion of these neurons to cause thermal hyperalgesia. Many previous studies have shown that PAR2 activation sensitizes TRPV1 (Amadesi et al., 2006; Amadesi et al., 2004; Dai et al., 2004), but many of these studies used SLIGRL, a PAR2-activating peptide that also stimulates Mrg receptors (Liu et al., 2011). The cellular basis of PAR2-mediated thermal hyperalgesia remains unresolved.

PAR2 has previously been implicated in itch, but this literature is also complicated by the non-specific nature of some widely used PAR2 pharmacological tools. For instance, SLIGRL-induced itch is mediated by an Mrg GPCR, and not by PAR2 (Liu et al., 2011). Nevertheless, some itch causing agents like cowhage activate PAR2 (Akiyama et al., 2015; Akiyama et al., 2009). Interestingly, our RNAscope and analysis of single cell RNA sequencing experiments clearly show that *F2rl1* mRNA is expressed in a population of DRG neurons that are known to be critical for itch behaviors in mice (Mishra and Hoon, 2013). Importantly, the contribution of this subset of neurons to nociception has not been clear. Because of these previous and current findings, we tested whether 2AT can cause pain or itch behaviors in mice. At 2AT doses that are specific for PAR2 (Boitano et al., 2014), we observed clear pain behaviors, consistent with our grimace findings. We did not observe itch behaviors. IL31, which acts via a receptor that is expressed by this same population of cells (Cevikbas et al., 2014), produced both itch and pain behaviors, indicating that activating these neurons is capable of driving both types of behavioral outcomes. These differential outcomes may be mediated by encoding at the level of the spinal cord, as it has recently been shown that burst firing in those itch circuits is required to drive itch behavior (Pagani et al., 2019; Petitjean et al., 2019). At high concentrations of 2AT, we noted both pain and itch behaviors, and the pattern of these behaviors were different in *F2rl1^-/-^* and WT mice. While we did not explore the mechanisms driving this difference experimentally, it may be explained by a complex pattern of recruitment of different populations of nociceptors because of the lack of specificity of the compound at concentrations above ~ 10 μM (Boitano et al., 2014). This pattern would necessarily be different in mice lacking PAR2. It could also be explained by differential innervation patterns for different types of afferents. In this regard, it has recently been shown that jugular neurons expressing an itch receptor, MrgprC11, signal bronchoconstriction and airway hyperresponsiveness (Han et al., 2018). The physiological outcome of PAR2 activation is likely dependent on the peripheral and central innervation target of the PAR2-expressing neuron.

Previous work from our group has demonstrated that activation of PAR2 leads to the development of a persistent pain state termed hyperalgesic priming (Tillu et al., 2015). This pain plasticity requires PAR2-mediated activation of ERK and downstream signaling to translation initiation factors that alter gene expression in nociceptors (Moy et al., 2017; Moy et al., 2018; Tillu et al., 2015). These PAR2-mediated effects could have been due to signaling in sensory neurons or other non-neuronal cells. Our current work demonstrates that this effect is mainly driven by nociceptor-expressed PAR2 because 2AT-evoked mechanical hypersensitivity and grimacing are gone with sensory neuron-specific deletion of the *F2rl1* gene and PAR2 activation activates ERK in this specific population of nociceptors. While we provide compelling evidence that most aspects of PAR2-mediated pain are due to PAR2 in nociceptors, it remains to be seen what tissues these nociceptors innervate. This is an important question to address in future studies. PAR2 has been implicated in gastro-intestinal pain (Cenac et al., 2007; Jimenez-Vargas et al., 2018), but few, if any, colonic sensory neurons express *F2rl1* mRNA (Hockley et al., 2018). Discovering the innervation pattern of this population of nociceptors will clarify which pain disorders are likely to benefit from PAR2 antagonist therapy.

---

**Supplementary Table 1 –.**
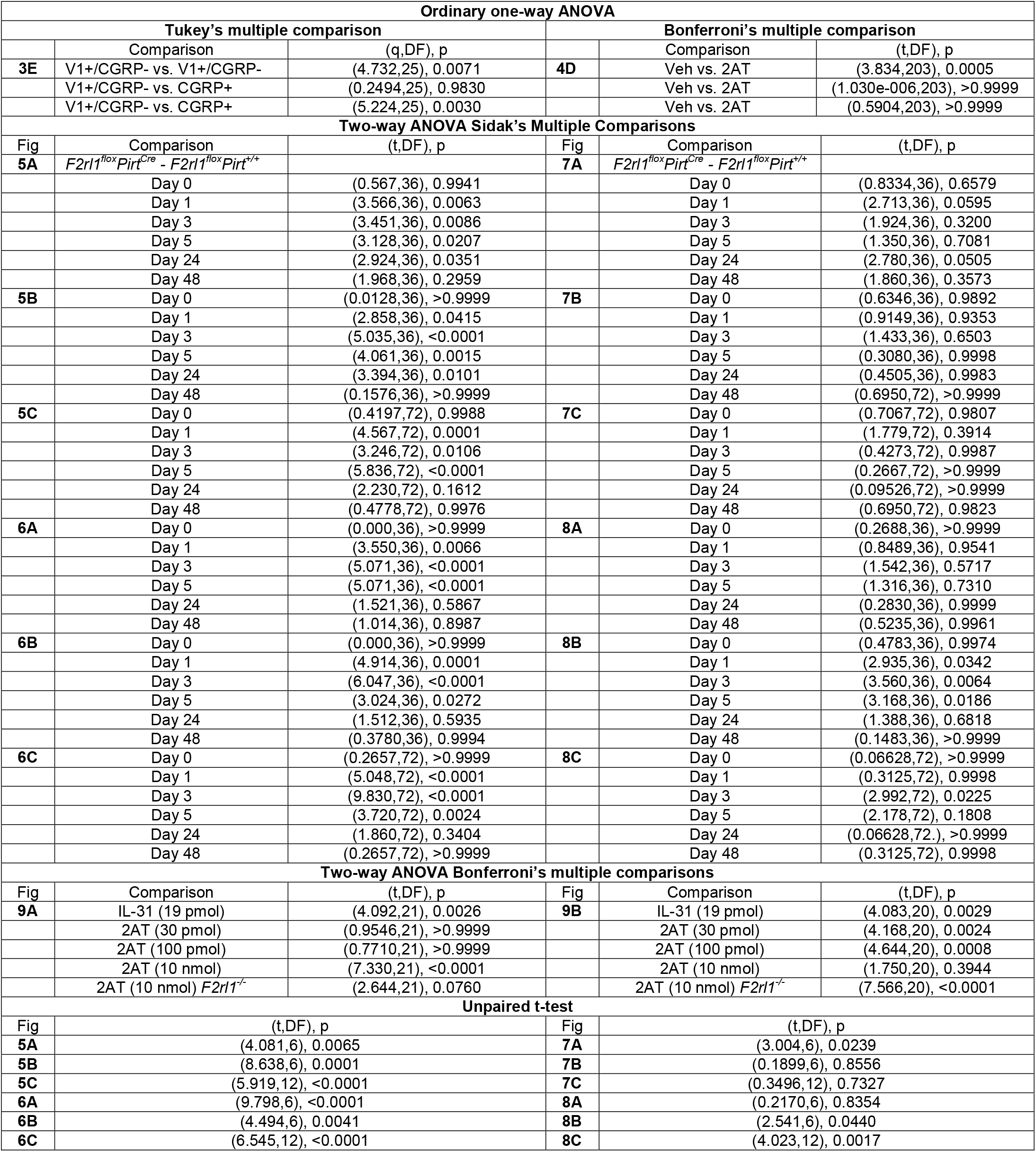
Ordinary one-way ANOVA Tukey’s and Bonferroni’s multiple comparisons, two-way ANOVA Sidak’s comparisons or unpaired t-test as indicated

**Supplementary Table 2 -.** Two-way ANOVA Dunnett’s comparisons

| Fig | Comparison | (q,DF), p | Comparison | (q,DF), p |
| --- | --- | --- | --- | --- |
|  | *F2rl1<sup>fllox</sup>Pirt<sup>Cre</sup>* |  | *F2rl1<sup>fllox</sup>Pirt<sup>+/+</sup>* |  |
| **5A** | Day 0 vs. Day 1 | (2.732,30), 0.0422 | Day 0 vs. Day 1 | (5.802,30), <0.0001 |
|  | Day 0 vs. Day 3 | (1.750,30), 0.2975 | Day 0 vs. Day 3 | (4.702,30), 0.0003 |
|  | Day 0 vs. Day 5 | (2.017,30), 0.1863 | Day 0 vs. Day 5 | (4.639,30), 0.0003 |
|  | Day 0 vs. Day 24 | (1.974,30), 0.2018 | Day 0 vs. Day 24 | (4.387,30), 0.0006 |
|  | Day 0 vs. Day 48 | (0.02667,30), >0.9999 | Day 0 vs. Day 48 | (1.408,30), 0.4959 |
| **5B** | Day 0 vs. Day 1 | (1.865,30), 0.2247 | Day 0 vs. Day 1 | (4.650,30), 0.0003 |
|  | Day 0 vs. Day 3 | (2.543,30), 0.0644 | Day 0 vs. Day 3 | (7.440,30), <0.0001 |
|  | Day 0 vs. Day 5 | (2.023,30), 0.1843 | Day 0 vs. Day 5 | (5.976,30), <0.0001 |
|  | Day 0 vs. Day 24 | (0.07999,30), 0.9999 | Day 0 vs. Day 24 | (3.225,30), 0.0130 |
|  | Day 0 vs. Day 48 | (0.3381,30), 0.9966 | Day 0 vs. Day 48 | (0.5035,30), 0.9815 |
| **5C** | Day 0 vs. Day 1 | (5.233,60), <0.0001 | Day 0 vs. Day 1 | (9.373,60), <0.0001 |
|  | Day 0 vs. Day 3 | (3.790,30), 0.0017 | Day 0 vs. Day 3 | (6.605,60), <0.0001 |
|  | Day 0 vs. Day 5 | (1.083,60), 0.7181 | Day 0 vs. Day 5 | (6.477,60), <0.0001 |
|  | Day 0 vs. Day 24 | (1.186,60), 0.6454 | Day 0 vs. Day 24 | (2.989,60), 0.0176 |
|  | Day 0 vs. Day 48 | (0.9787,60), 0.7879 | Day 0 vs. Day 48 | (0.08473,60), 0.9999 |
| **6A** | Day 0 vs. Day 1 | (0.000,30), >0.9999 | Day 0 vs. Day 1 | (3.337,30), 0.0098 |
|  | Day 0 vs. Day 3 | (0.476,30), 0.9854 | Day 0 vs. Day 3 | (5.244,30), <0.0001 |
|  | Day 0 vs. Day 5 | (0.476,30), 0.9854 | Day 0 vs. Day 5 | (5.244,30), <0.0001 |
|  | Day 0 vs. Day 24 | (1.059e-015,30), >0.9999 | Day 0 vs. Day 24 | (1.430,30), 0.4812 |
|  | Day 0 vs. Day 48 | (0.000,30), >0.9999 | Day 0 vs. Day 48 | (0.9535,30), 0.8035 |
| **6B** | Day 0 vs. Day 1 | (1.348,30), 0.5361 | Day 0 vs. Day 1 | (7.187,30), <0.0001 |
|  | Day 0 vs. Day 3 | (0.8984,30), 0.8362 | Day 0 vs. Day 3 | (8.086,30), <0.0001 |
|  | Day 0 vs. Day 5 | (0.8984,30), 0.8362 | Day 0 vs. Day 5 | (4.492,30), 0.0005 |
|  | Day 0 vs. Day 24 | (4.987e-016,30), >0.9999 | Day 0 vs. Day 24 | (1.797,30), 0.2751 |
|  | Day 0 vs. Day 48 | (9.975e-016,30), >0.9999 | Day 0 vs. Day 48 | (0.4492,3), 0.9888 |
| **6C** | Day 0 vs. Day 1 | (0.8478,60), 0.8653 | Day 0 vs. Day 1 | (6.500,60), <0.0001 |
|  | Day 0 vs. Day 3 | (0.5652,60), 0.9709 | Day 0 vs. Day 3 | (11.30,60), <0.0001 |
|  | Day 0 vs. Day 5 | (0.5652,60), 0.9709 | Day 0 vs. Day 5 | (4.804,60), <0.0001 |
|  | Day 0 vs. Day 24 | (1.098e-015,60)>0.9999 | Day 0 vs. Day 24 | (2.261,60), 0.1051 |
|  | Day 0 vs. Day 48 | (0.2826,60), 0.9984 | Day 0 vs. Day 48 | (0.2826,60), 0.9984 |
| **7A** | Day 0 vs. Day 1 | (0.5954,30), 0.9630 | Day 0 vs. Day 1 | (2.471,30), 0.0752 |
|  | Day 0 vs. Day 3 | (1.415,30), 0.4911 | Day 0 vs. Day 3 | (2.503,30), 0.0701 |
|  | Day 0 vs. Day 5 | (0.06533,30), >0.9999 | Day 0 vs. Day 5 | (0.5806,30), 0.9665 |
|  | Day 0 vs. Day 24 | (1.512,30), 0.4296 | Day 0 vs. Day 24 | (3.454,30), 0.0073 |
|  | Day 0 vs. Day 48 | (0.1470,30), 0.9998 | Day 0 vs. Day 48 | (1.172,30), 0.6582 |
| **7B** | Day 0 vs. Day 1 | (2.212,30), 0.1281 | Day 0 vs. Day 1 | (2.515,30), 0.0684 |
|  | Day 0 vs. Day 3 | (1.758,30), 0.2936 | Day 0 vs. Day 3 | (2.618,30), 0.0545 |
|  | Day 0 vs. Day 5 | (2.981,30), 0.0236 | Day 0 vs. Day 5 | (1.964,30), 0.2054 |
|  | Day 0 vs. Day 24 | (2.537,30), 0.0652 | Day 0 vs. Day 24 | (2.339,30), 0.0992 |
|  | Day 0 vs. Day 48 | (2.997,30), 0.0227 | Day 0 vs. Day 48 | (2.442,30), 0.0799 |
| **7C** | Day 0 vs. Day 1 | (3.373,60), 0.0059 | Day 0 vs. Day 1 | (4.518,60), 0.0001 |
|  | Day 0 vs. Day 3 | (4.138,60), 0.0006 | Day 0 vs. Day 3 | (2.928,60), 0.0207 |
|  | Day 0 vs. Day 5 | (3.691,60), 0.0023 | Day 0 vs. Day 5 | (3.221,60), 0.0092 |
|  | Day 0 vs. Day 24 | (3.757,60), 0.0018 | Day 0 vs. Day 24 | (3.104,60), 0.0128 |
|  | Day 0 vs. Day 48 | (3.877,60), 0.0013 | Day 0 vs. Day 48 | (3.864,60), 0.0013 |
| **8A** | Day 0 vs. Day 1 | (0.3268,30), 0.9970 | Day 0 vs. Day 1 | (0.2558,30), 0.9987 |
|  | Day 0 vs. Day 3 | (1.080,30), 0.7213 | Day 0 vs. Day 3 | (0.1989,30), 0.9997 |
|  | Day 0 vs. Day 5 | (0.04263,30), >0.9999 | Day 0 vs. Day 5 | (1.549,30), 0.4070 |
|  | Day 0 vs. Day 24 | (0.3268,30), 0.9970 | Day 0 vs. Day 24 | (0.3126,30), 0.9977 |
|  | Day 0 vs. Day 48 | (0.2984,30), 0.9982 | Day 0 vs. Day 48 | (0.04263,30), >0.9999 |
| **8B** | Day 0 vs. Day 1 | (0.5990,30), 0.9620 | Day 0 vs. Day 1 | (2.409,30), 0.0857 |
|  | Day 0 vs. Day 3 | (0.3777,30), 0.9949 | Day 0 vs. Day 3 | (3.399,30), 0.0084 |
|  | Day 0 vs. Day 5 | (0.8595,30), 0.8578 | Day 0 vs. Day 5 | (2.4350,30), 0.0811 |
|  | Day 0 vs. Day 24 | (2.513,30), 0.0686 | Day 0 vs. Day 24 | (3.620,30), 0.0048 |
|  | Day 0 vs. Day 48 | (0.3386,30), 0.9966 | Day 0 vs. Day 48 | (0.4428,30), 0.9895 |
| **8C** | Day 0 vs. Day 1 | (2.463,60), 0.0666 | Day 0 vs. Day 1 | (2.821,60), 0.0274 |
|  | Day 0 vs. Day 3 | (1.495,60), 0.4315 | Day 0 vs. Day 3 | (4.338,60), 0.0002 |
|  | Day 0 vs. Day 5 | (1.352,60), 0.5273 | Day 0 vs. Day 5 | (3.474,60), 0.0044 |
|  | Day 0 vs. Day 24 | (0.9581,60), 0.8010 | Day 0 vs. Day 24 | (1.084,60), 0.7177 |
|  | Day 0 vs. Day 48 | (0.2776,60), 0.9985 | Day 0 vs. Day 48 | (0.08059,60), 0.9999 |

## ACKNOWLEDGEMENTS

This work was supported by NIH grants R01NS065926 (TJP) R01NS102161 (TJP and ANA) and R01NS098826 (TJP, SB, JV and GD)

